# Structural Analysis of Wild-type and Val120Thr Mutant *Candida boidinii* Formate Dehydrogenase by X-ray Crystallography

**DOI:** 10.1101/2022.12.25.521900

**Authors:** Mehmet Gul, Busra Yuksel, Huri Bulut, Hasan DeMirci

## Abstract

Candida boidinii NAD^+^-dependent formate dehydrogenase (CbFDH) has gained significant attention for its potential applications in the production of biofuels and various industrial chemicals from inorganic carbon dioxide. The present study reports the atomic X-ray crystal structures of the wild-type CbFDH at cryogenic and ambient temperatures as well as Val120Thr mutant at cryogenic temperature determined at the Turkish Light Source “Turkish DeLight”. The structures reveal new hydrogen bonds between Thr120 and water molecules in the mutant CbFDH’s active site, suggesting increased stability of the active site and more efficient electron transfer during the reaction. Further experimental data is needed to test these hypotheses. Collectively, our findings provide invaluable insights into future protein engineering efforts that could potentially enhance the efficiency and effectiveness of CbFDH.

## 1. Introduction

Candida boidinii NAD^+^ -dependent formate dehydrogenase (CbFDH) is a homodimeric 82 KDa enzyme complex that catalyzes the reversible conversion between formate and carbon dioxide (Nielsen et al. 2019). It catalyzes the reversible conversion of formate and carbon dioxide using NAD^+^/NADH coenzymes for electron transfer (Niks & Hille 2018). Many prokaryotic and eukaryotic organisms express NAD^+^-dependent FDH enzymes with similar structural and kinetic properties (Popov & Lamzin 1994). However, many studies have focused on the Candida boidinii NAD^+^-dependent FDH (CbFDH) due to its remarkable stability and activity (Bulut et al. 2021). Formate dehydrogenase was first discovered in the 1950s and recently gained great attention as a research topic due to its potential for formate molecule production from inorganic carbon dioxide. Formate is a valuable precursor molecule used in the synthesis of industrial chemicals such as formaldehyde, which has numerous applications in the production of resins, plastics, and disinfectants (“Formaldehyde,” 2016; Murashima et al. 2022; Powers 1953). Other chemicals that can be generated from formate include formic acid and oxalic acid, both of which have a range of important industrial applications (Schuler et al. 2021). Moreover, the formate molecule has the potential as an alternative biofuel since it can be converted into hydrogen gas by electrolysis (Eppinger & Huang 2017). Overall, formate’s versatility in industrial applications has made the FDH enzyme a central research topic.

Formate dehydrogenase also emerges a highly promising candidate for carbon capture efforts to counteract global climate change. The enzyme’s inherent capability to catalyze the conversion of CO_2_ to formate holds significant potential for reducing atmospheric CO_2_ levels (Calzadiaz-Ramirez & Meyer, 2022; Choe et al., 2014; Miller et al., 2019). Nonetheless, to optimize the efficiency and robustness in CO_2_ conversion, studies that focus on engineering FDH are necessary. These involve increasing its thermostability to endure harsh conditions and enhancing enzymatic activity for the reverse reaction, making it an even more suitable option for carbon capture strategies (Calzadiaz-Ramirez & Meyer, 2022). In order to enhance stability and activity, researchers have made significant progress by identifying and using enzymes from organisms that live in extreme environments. While Candida boidinii might not be considered an extremophile, these studies have shown the potential of extremophiles as valuable genetic resources. Extremophiles have provided insights into making enzymes stable in different industrial and environmental situations. The stability-enhancing adaptations observed in extremophilic enzymes may inspire protein engineers to employ similar strategies to improve enzymes like *Cb*FDH, even when the host organism is not an extremophile (Chen & Jiang, 2018; Dumorne et al., 2017; Ye et al., 2023).

Proteins’ industrial and therapeutic applications are significantly impacted by their limited activity and stability under various industrial conditions. Therefore, protein engineering studies are crucial to generate mutant proteins to overcome the limitations and improve protein function. Protein engineering studies with the help of computational biology tools allow us to identify and mutate the amino acid residues that may improve the thermostability of proteins. Thermostable mutants are suitable for industrial biocatalysis due to their increased resistance to high temperatures. These mutations include single or multiple amino acid alterations, insertions, or deletions for greater numbers of hydrogen bonds, disulfide bonds, or salt bridges to increase thermostability (Modarres et al. 2016). In addition, protein engineering approaches are also focused on creating proteins with higher enzymatic activity. The more active mutant proteins can increase the rate and efficiency of the reaction. Similar to stability, protein engineering studies are used to identify and modify the amino acids that can improve activity, specificity and substrate affinity. Site-directed mutagenesis, random mutagenesis, and DNA shuffling methods are widely used to generate highly active mutant proteins (Li et al. 2020). Within *Cb*FDH, the 120^th^ residue, Val120, is a potential mutation site for studies aiming to create a more active enzyme variant as it is located in the catalysis and substrate-binding site (Supplementary Figure 1). Although it is unclear whether Val120 is directly involved in the reaction, mutating it into a more polar residue, such as serine, results in enhanced catalytic activity. It might be due to changes in charge distribution within the active site of the enzyme (Jiang et al., 2016).

High-resolution structural studies are crucial for understanding the function, underpinnings of the catalytic mechanism, and other potential applications of dehydrogenase enzymes like *Cb*FDH. Previous studies have contributed to our knowledge of the structural and functional features of *Cb*FDH However, there are still gaps in our understanding of its function and potential modifications for further enhancing activity and thermal stability. Here, we present high-resolution crystal structures of the apo *Cb*FDH determined at cryogenic and ambient temperature, as well as a highly active *Cb*FDH Val120Thr mutant at cryogenic temperature. Our findings, combined with existing knowledge of the enzyme’s structure and function, will drive further research on the potential industrial applications of *Cb*FDH.

## 2 Results

### 2.1 Wild-type apo CbFDH structure is determined at 1.4 Å resolution at cryogenic temperature

We determined the crystal structure of wild-type apo *Cb*FDH at 1.4 Å resolution (PDB ID: 8HTY) with overall 98.82% completeness at cryogenic temperature at the Turkish Light Source “Turkish DeLight” (Istanbul, Türkiye). The determined structure consists of two homodimers of *Cb*FDH (Figure 1). The Ramachandran plot indicates that 98.01% of the residues were in the favored regions while 1.99% were in the allowed regions with no outliers. The data collection and refinement statistics are given in Table 1.

**Figure 1.**
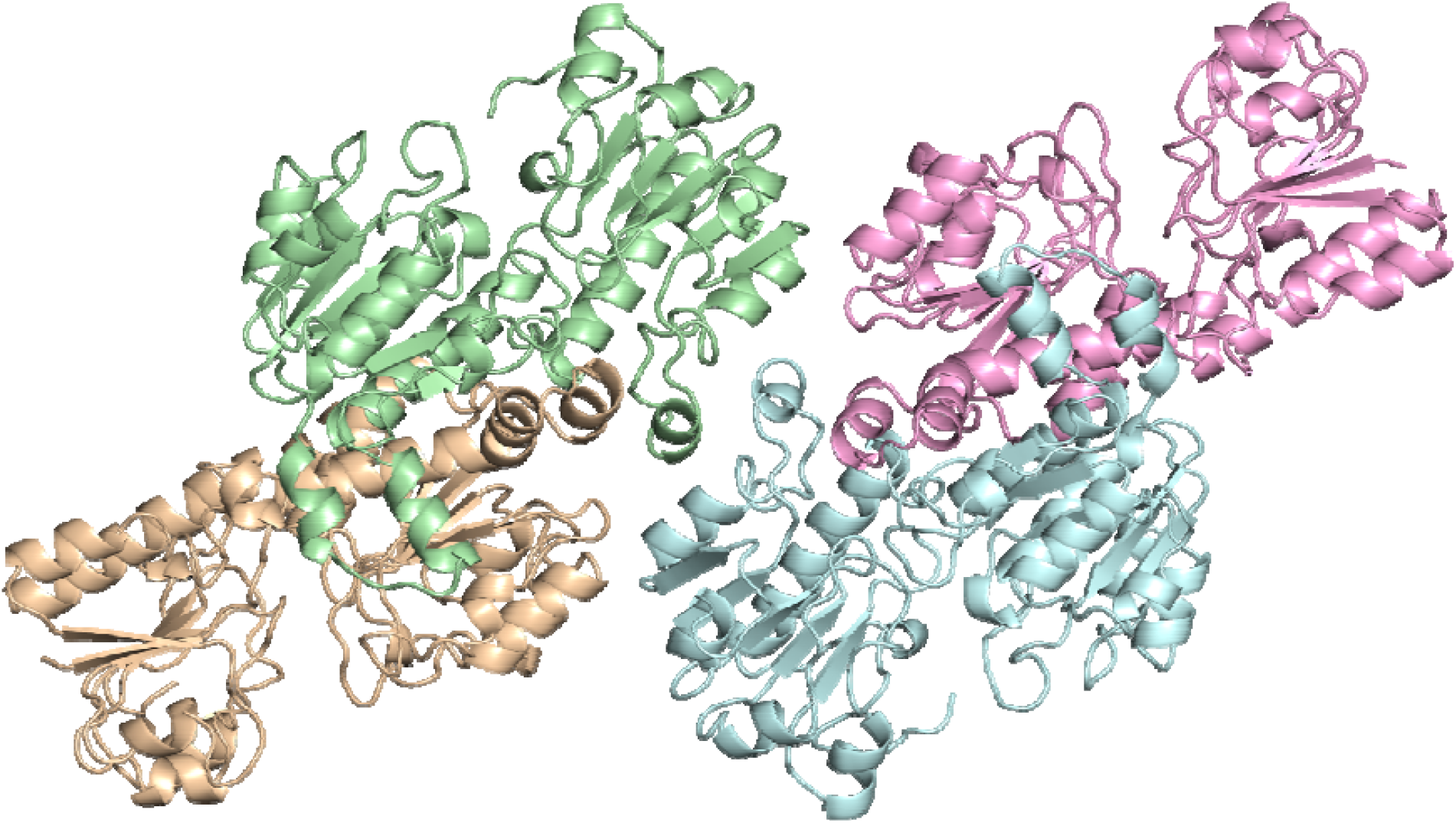
1.4 Å apo wild-type *Cb*FDH structure as two homodimers. Four monomer molecules are shown in different colors. Monomers in pale green and wheat form the first dimer, while monomers in pink and pale cyan form the second dimer.

**Table 1.** Data collection and refinement statistics. The highest resolution shell is shown in parentheses. CT: cryogenic temperature; AT: ambient temperature.

| Dataset | Wt-*CbFDH*-CT | Wt-*CbFDH*-AT | Val120Thr-*CbFDH*-CT |
| --- | --- | --- | --- |

| PDB ID | 8HTY | 8IQ7 | 8IVJ |
| --- | --- | --- | --- |
| **Data collection** |  |  |  |
| X-ray source | *Turkish DeLight* | *Turkish DeLight* | *Turkish DeLight* |
| Wavelength (Å) | 1.54 | 1.54 | 1.54 |
| Temperature (K) | 100 | 298 | 100 |
| Space group | P 1 | P 1 | P 1 |
| Cell dimensions |  |  |  |
| *a*, *b*, *c* (Å) | 54.1, 69.0, 109.8 | 54.7, 69.5, 110.8 | 53.7, 68.6, 109.4 |
| $\alpha$, $\beta$, $\gamma$ (°) | 78.1, 89.5, 81.1 | 77.9, 89.1, 81.2 | 78.0, 89.6, 81.4 |
| Resolution (Å) | 24.75-1.39 (1.51-1.39) | 30.73-2.10 (2.15-2.10) | 25.42-1.90 (1.94-1.90) |
| *CC* (½) | 0.999 (0.340) | 0.916 (0.196) | 0.622 (0.535) |
| *CC* * | 1.000 (0.708) | 0.978 (0.372) | 0.876 (0.835) |
| *I* / σ*I* | 11.74 (0.73) | 7.90 (0.60) | 3.00 (0.43) |
| Completeness (%) | 98.82 (86.0) | 85.90 (77.30) | 99.7 (99.5) |
| Redundancy | 151.46 (148.36) | 7.50 (3.30) | 7.30 (4.60) |
| **Refinement** |  |  |  |
| Resolution (Å) | 21.83-1.40 (1.43-1.40) | 30.70-2.10 (2.15-2.10) | 24.30-1.90 (1.95-1.90) |
| No. reflections | 301717 (29123) | 67181 (4741) | 96104 (7897) |
| *R*<sub>work</sub> / *R*<sub>free</sub> | 0.167/0.190<br>(0.380/0.372) | 0.2433/0.2716<br>(0.4364/0.4810) | 0.2469/ 0.2712<br>(0.4052/ 0.4226) |
| No. atoms |  |  |  |
| Protein | 22225 | 22154 | 22275 |
| Ligand / Ion / Water | 2249 | 393 | 1459 |
| *B*-factors |  |  |  |
| Protein | 22.94 | 18.89 | 22.47 |
| Ligand / Ion / Water | 35.07 | 17.02 | 26.27 |
| Coordinate errors | 0.18 | 0.38 | 0.30 |
| R.m.s deviations |  |  |  |
| Bond lengths (Å) | 0.015 | 0.002 | 0.002 |
| Bond angles (°) | 1.422 | 0.545 | 0.568 |
| Ramachandran plot |  |  |  |
| Favored (%) | 97.4 | 97.66 | 98.02 |
| Allowed (%) | 2.6 | 2.34 | 1.98 |
| Disallowed (%) | - | - | - |

The resulting final electron density map was superior-quality and reveals the structural features such as coordinated water molecules, and side chains of the *Cb*FDH molecule in great details. The electron density remains intact and visible at 3 sigma levels (Figure 2).

**Figure 2.**
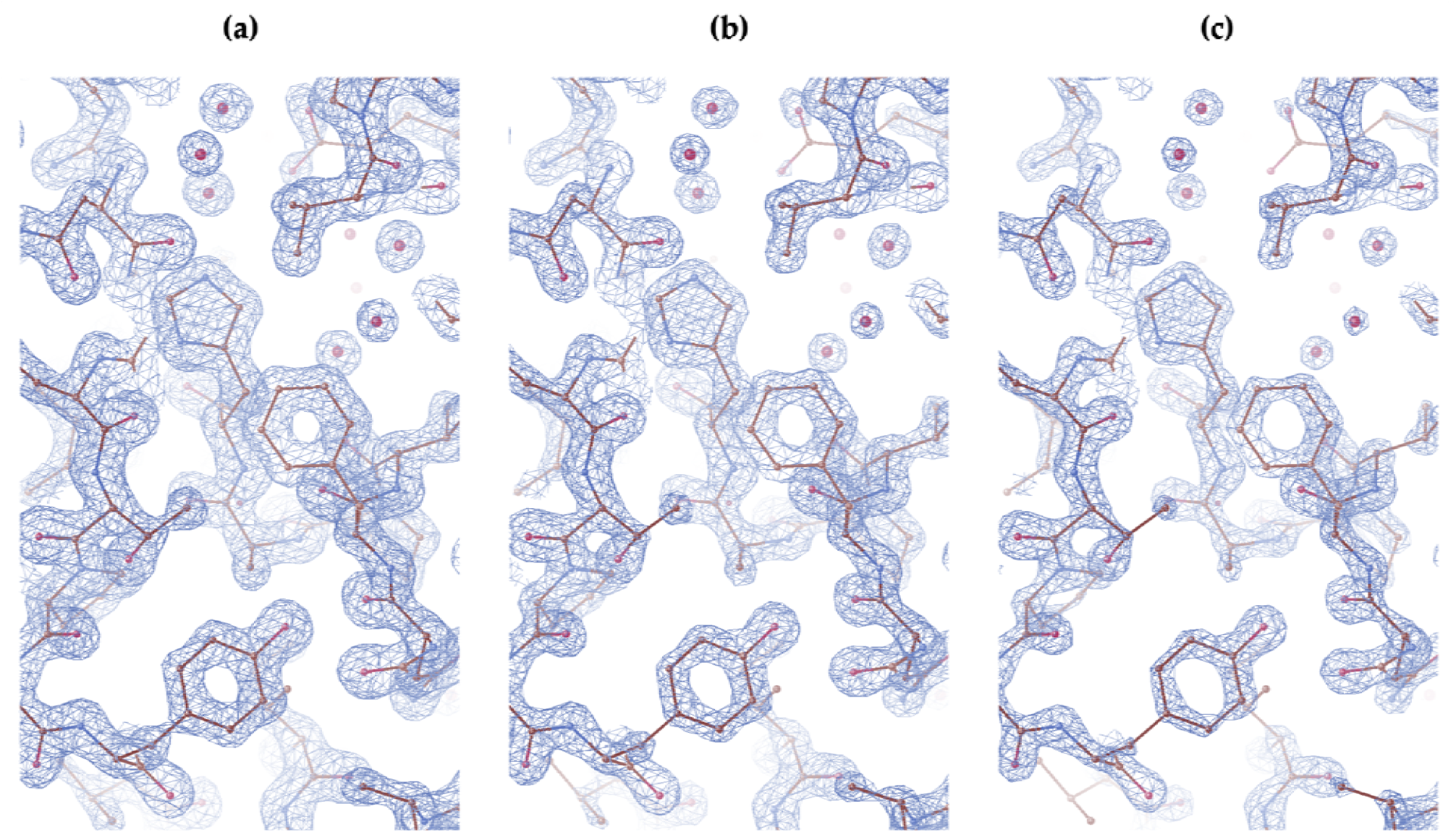
2Fo-Fc map is colored in blue and contoured at (a) 2.0 σ, (b) 2.5 σ, and (c) 3.0 σ level.*Cb*FDH is shown in stick representation.

We superposed the two CbFDH homodimers in our crystal structure to compare their similarities (Supplementary Figure 2). The two homodimers aligned with an RMSD score of 0.352. Particularly, the dimerization regions within the *Cb*FDH homodimers seem almost identical (Supplementary Figure 2c) while some minor conformational changes are observed in regions farther from the central dimerization core domain (Supplementary Figure 2b,d).

### 2.2 Two wild-type apo structures align with some minor differences

Our wild-type apo *Cb*FDH structure crystallized in the same space group and unit cell as the previous structure (PDB ID: 5DNA) (Guo et al., 2016). When we compare the two structures the overall structures align very well with an RMSD of 0.266 Å as expected (Figure 3).

**Figure 3.**
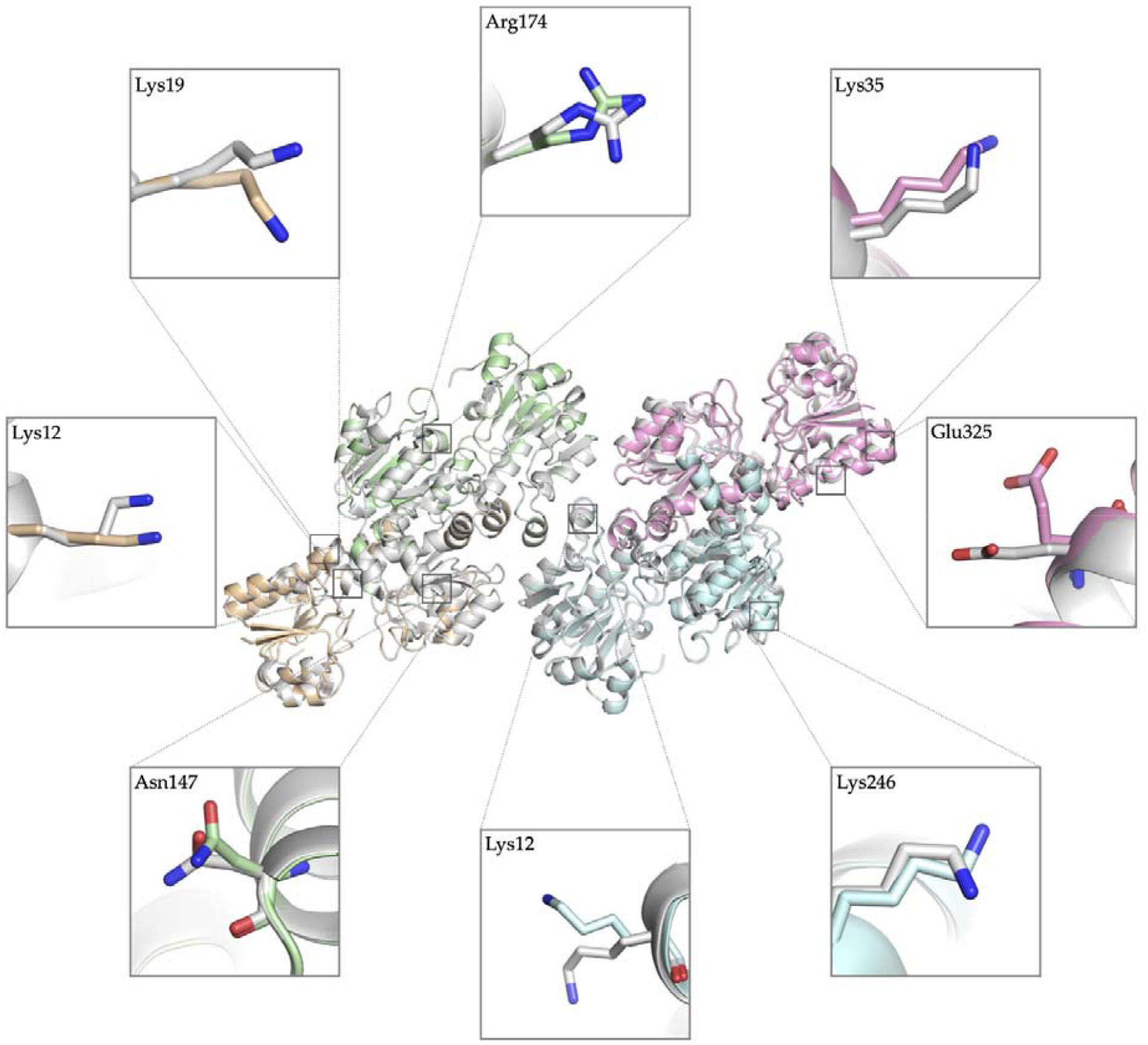
Superposition of two apo wild-type *Cb*FDH structures. Our 1.4 Å structure is shown in four distinct colors indicating each chain; A: pale green, B: wheat, C: pink, D: pale cyan. 5DNA is colored gray. Conformational differences in side chains are shown in boxes in stick representation.

Differences are mostly observed in the residues that directly exposed to the solvent (Figure 3). In addition, the previous apo CbFDH structure lacks the residues 15-18 (Ala15, Asp16, Glu17, Glu18), which are part of a loop in chain C, while our structure has a well-defined electron density for them (Supplementary Figure 3).

### 2.3 Wild-type apo *Cb*FDH structure is determined at 2.1 Å resolution at ambient temperature

The crystal structure of wild-type apo CbFDH was determined at at the Turkish Light Source “Turkish DeLight” (Istanbul, Türkiye) at 2.1 Å resolution (PDB ID: 8IQ7) with 85.9% completeness at ambient temperature. Similar to the cryogenic structure, the structure consist of two homodimers (Figure 4). 97.66% of the residues were in the favored regions while 2.34% were in the allowed regions with no outliers according to the Ramachandran statistics. The data collection and refinement statistics are given in Table 1.

**Figure 4.**
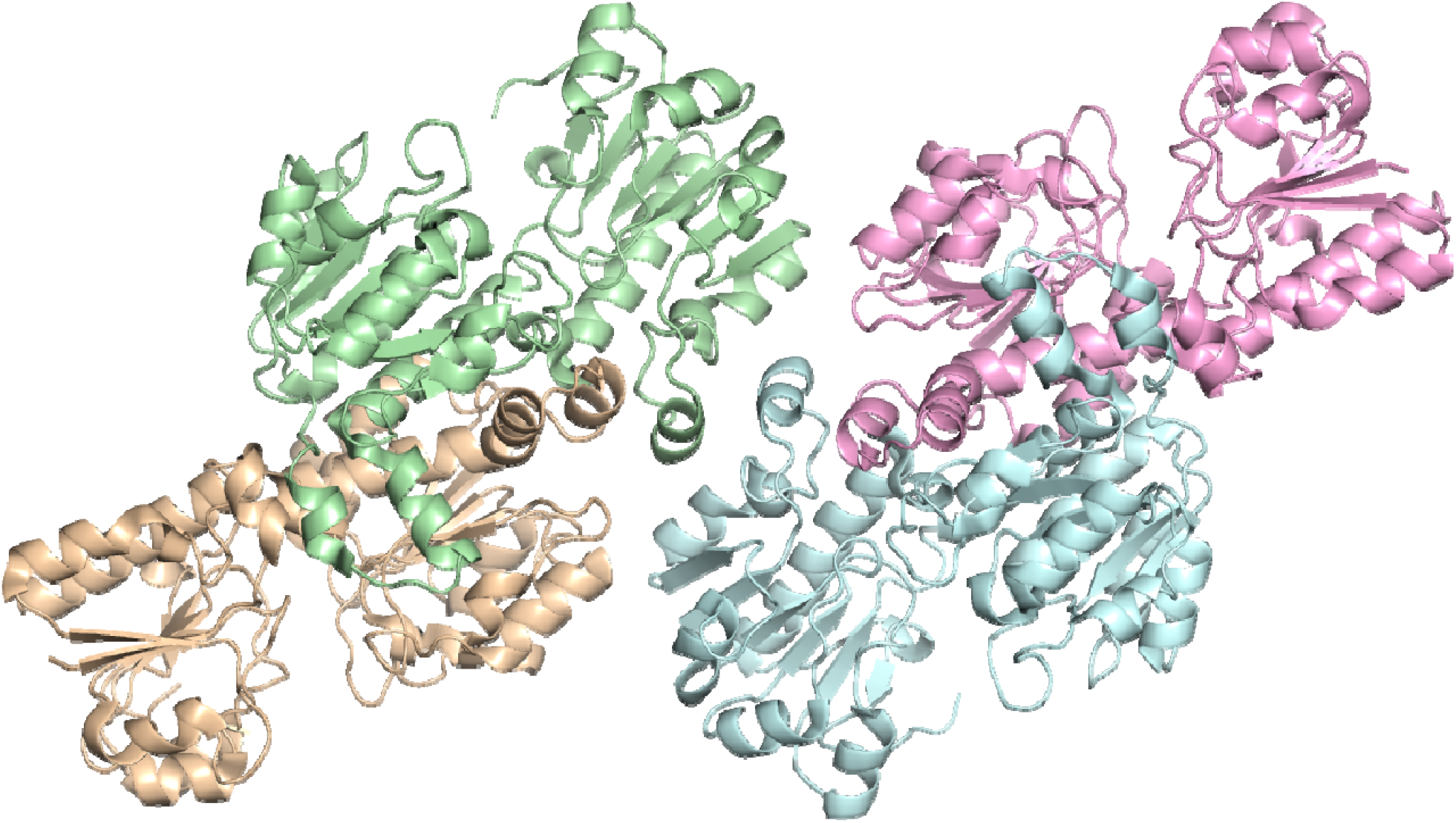
Wild-type apo *Cb*FDH structure at 2.1 Å resolution at ambient temperature.

Cryogenic and ambient apo *Cb*FDH structures possess slight differences Our cryogenic (PDB ID: 8HTY) and ambient (PDB ID: 8IQ7) *Cb*FDH structures are superposed with an RMSD of 0.554, suggesting slight differences between the two structures. These differences are mostly in the flexible loop regions and amino acid side chains (Figure 5).

**Figure 5.**
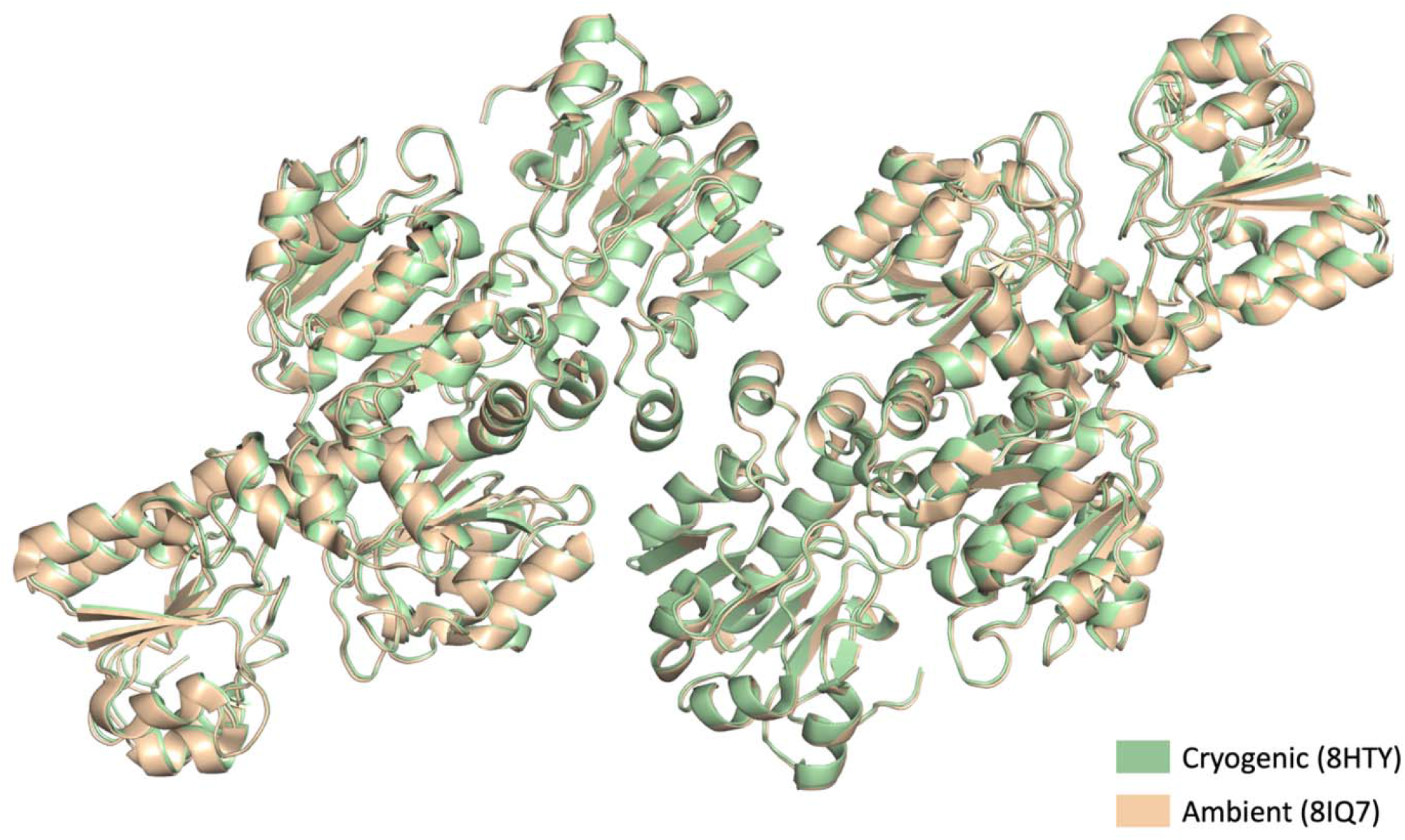
Superposition of cryogenic and ambient wild-type *Cb*FDH structures.

We also compared the number of water molecules identified within our cryogenic and ambient temperature structures. The cryogenic temperature structure exhibited a significantly higher number of identified water molecules, with 2179 waters resolved, compared to the ambient temperature structure with only 393 water molecules.

### 2.5 Mutation from valine to threonine significantly enhances *Cb*FDH activity

Kinetic assays were performed to assess the changes in *Cb*FDH activity due to mutation from valine to threonine at the 120th residue. The K_m_, k_cat_, and k_cat_/K_m_ values calculated for formate are shown in Table 2. The lower K_m_ value for formate in Val120Thr mutant indicates approximately 1.5-fold stronger affinity for substrate binding while higher k_cat_ shows the higher turnover number (6.57-fold) compared to the wild-type. The overall catalytic efficiency (k_cat_/K_m_) is found approximately ten times greater in the mutant *Cb*FDH.

**Table 2.**
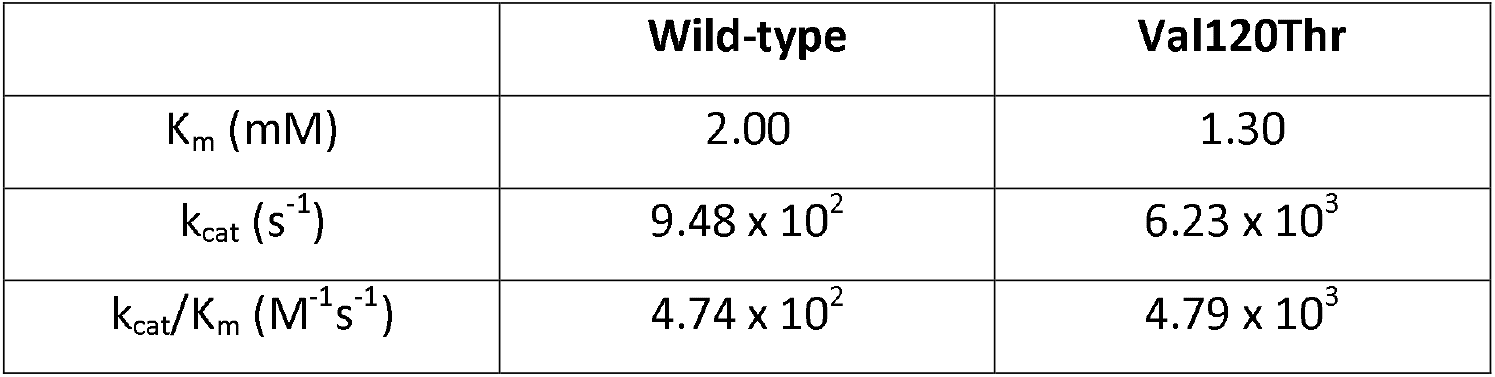
Kinetic assay results for wild-type and Val120Thr *Cb*FDH.

|  | Wild-type | Val120Thr |
| --- | --- | --- |
| $K_m$ (mM) | 2.00 | 1.30 |
| $k_{cat}$ ($s^{-1}$) | $9.48 \times 10^2$ | $6.23 \times 10^3$ |
| $k_{cat}/K_m$ ($M^{-1}s^{-1}$) | $4.74 \times 10^2$ | $4.79 \times 10^3$ |

### 2.6 *Cb*FDH Val120Thr mutant structure reveals new hydrogen bondings in the active site

We determined the structure of our novel Val120Thr mutant *Cb*FDH at 1.9 Å resolution at cryogenic temperature (PDB ID: 8IVJ). The structure is composed of two homodimers with 99.7% completeness. The Ramachandran statistics indicate that 98.02% of the residues are in the favored and 1.98% in the allowed regions with no outliers. Table 1 shows the data collection and refinement statistics.

Superposition of the wild-type *Cb*FDH and Val120Thr mutant (Supplementary Figure 4a) demonstrated that the overall structure does not change significantly with the single mutation (RMSD: 0.270). However, a closer look to the mutation site revealed new hydrogen bonds that does not exist in the wild-type CbFDH between the 120^th^ residue threonine and three water molecules (Figure 6; Supplementary Figure 4b-f).

**Figure 6.**
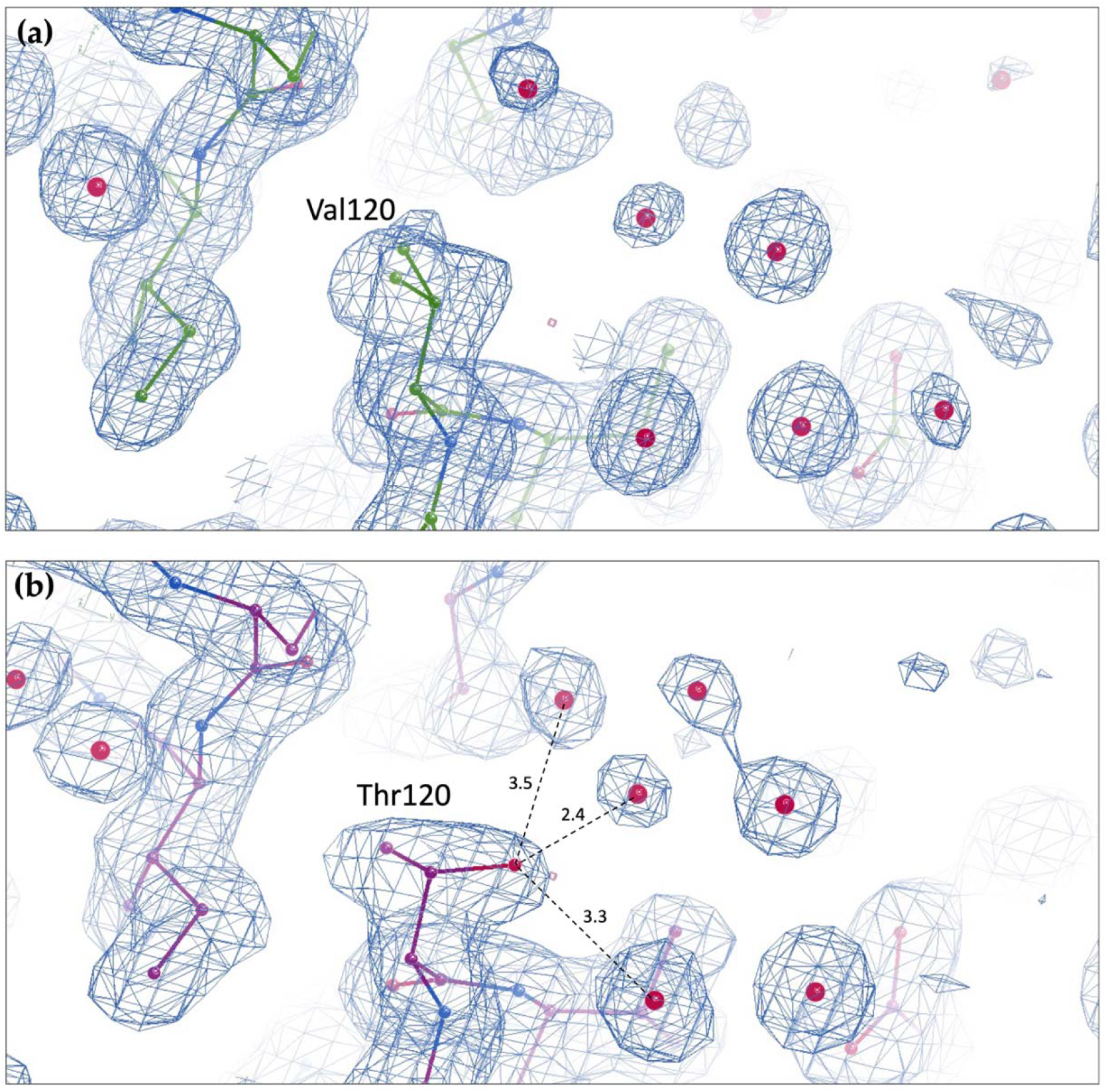
2Fo-Fc simulated annealing-omit map of the 120th residue of *Cb*FDH, shown in blue and contoured at 1.0 σ level (A. T. Brünger et al., 1998; Hodel, Kim, & Brünger, 1992). (a) Val120 of the cryogenic wild type *Cb*FDH structure (8HTY). (b) Thr120 of the cryogenic Val120Thr mutant *Cb*FDH (8IVJ). Hydrogen bonding distances in Angstroms are indicated with dashed lines.

## 3. Discussion

Our 1.4 Å-resolution apo CbFDH structure provides structural details of the enzyme in its homodimeric form. The superior-quality electron density map reveals atomic details of the amino acid residues and water molecules. This high-resolution structure revealed a high degree of similarity between the two subunits, with only minor differences observed in the more flexible regions outside of the dimerization core domain region. These minor changes are expected since they can move more freely as they do not participate in dimer formation.

While the two homodimers in our structure were highly identical, a comparison with a previously published apo structure, consisting of two homodimers as well, (PDB ID: 5DNA) at a 1.75 Å resolution revealed several minor differences, particularly in the side chains of solvent exposed charged residues. Furthermore, our structure also showed improved electron density for flexible loop residues 15-18 (Ala15, Asp16, Glu17, Glu18) in chain C that were not well defined in the previous structure.

Studying protein structures at ambient temperature offers valuable insights into their natural conformational dynamics, active site flexibility, functional mechanisms, and binding behavior. This approach may capture a broader range of conformational states, especially for proteins that are sensitive to cryogenic temperatures (Fischer, 2021; Fraser et al., 2011). By combining insights from both cryogenic and ambient temperature structures, we may gain a more complete understanding of protein structures and their dynamics. Therefore, in addition to the structure obtained at cryogenic temperature, we also determined the structure of the wild-type CbFDH at ambient temperature. The comparison between the wild-type structures determined at cryogenic and ambient temperature showed that the structure did not significantly change due to temperature except for flexible loops and large amino acid side chains. The reason might be that the flash-freezing in liquid nitrogen captured the protein at its most stable state in solution, which we also observed in the ambient temperature structure. Still, the main difference between the two structures is most likely in the flexibility or mobility. However, the resolution difference may affect the comparison of the b-factor. Thus, further analysis is required to fully understand the impact of the temperature on the structure.

The significant difference in the number of water molecules identified within the cryogenic and ambient temperature raises intriguing questions about the dynamic interactions between protein and water under different environmental conditions. The higher resolution achieved at cryogenic temperatures significantly improves the accuracy of identifying water molecules compared to the ambient temperature structures with lower resolution. Another reason for different numbers of identified water molecules could be the differences in water accessibility and occupancy between the two conditions. At cryogenic temperatures, water molecules might be frozen at specific binding sites, leading to the stabilization of interactions that are transient at ambient temperature (Nakasako, 2004). This could explain why cryogenic temperature structures allow for the identification of more water molecules compared to ambient temperature structures. These findings emphasize the importance of investigating temperature effects as the protein’s microenvironment exhibits high complexity with changes in conformation and water interactions due to environmental factors. Further studies, such as molecular dynamics simulations and spectroscopic techniques, can provide further insights into protein-water interactions and water molecule behavior.

According to the kinetic assay results, the mutant CbFDH has a lower K_m_ value, indicating a higher affinity to formate, and a higher kcat value, indicating a faster conversion to the product. Overall, the catalytic efficiency has been increased approximately ten times compared to the wild-type enzyme. The reason for the increase in substrate affinity may be that threonine is more polar than valine, so that it can form hydrogen bonds with neighboring amino acids or water molecules, resulting in higher stability in the substrate binding region. In addition, it may allow for an easier substrate binding by providing a larger space for formate to enter as threonine has a smaller side chain compared to valine.

In order to observe the effect of the Val120Thr mutation, we determined the structure of the mutant *Cb*FDH at 1.9 Å resolution and compared it with our wild-type structure. When the structures were superimposed, we did not observed major changes in the structure as expected. In the mutation site, nonetheless, we observed three new hydrogen bonds between Thr120 and water molecules in the active site. These new hydrogen bonds may help stabilize the active site, leading to greater activity. Also, the newly formed hydrogen bond may help positioning the substrate or coenzyme molecules in the active site and contribute to a more efficient electron transfer process during the reaction. Additionally, the hydroxyl group of the threonine side chain could affect the binding of the coenzyme NAD^+^ by interacting with it or other amino acids in the active site. In order to test these hypotheses, further experimental data is needed, including co-crystallization of the Val120Thr mutant CbFDH with NAD^+^ and formate/azide.

The high-resolution structures of the apo *Cb*FDH presented in this study provide insights into the enzyme’s structural details. Moreover, the Val120Thr mutation in the active site of the enzyme enhanced enzymatic activity by approximately ten times compared to the wild-type enzyme. The structural changes observed between the wild-type and mutant *Cb*FDH emphasize the significance of structural analysis in understanding the effects of mutations on enzyme activity. Further research on the mutant enzyme with its substrate and coenzyme is required to fully understand the underlying mechanism of the mutation. Overall, the findings of this study can potentially contribute to the development of enzymes with higher efficiency and specificity for a range of applications in biotechnology, industry, and medicine.

## 4. Materials and Methods

### 4.1 Transformation, expression, purification, and crystallization

Full-length wild-type and Val120Thr mutant CbFDH genes were cloned into pET-23a (+) bacterial expression vector with a C-terminal hexahistidine-tag as previously described by Bulut et al. (2021). The plasmids were transformed into the competent E. coli BL21 Rosetta-2 strain. Transformed bacterial cells were grown in 6 L LB medium containing 100 Μg/ml ampicillin and 35 Μl/ml chloramphenicol at 37 °C. Once OD600 reached to 0.7-0.8 range, the protein expression was induced by addition of β-D-1-thiogalactopyranoside (IPTG) to a final concentration of 0.4 mM for 3-4 hours at 37 °C. Cells were then harvested using Beckman Allegra 15R desktop centrifuge at 4 °C at 3500 rpm for 30 minutes. Cell pellets were stored at - 80 °C until purification.

The frozen cell pellets were dissolved in ice cold lysis buffer that contains 500 mM NaCl, 50 mM Tris-HCl pH 7.5, 5% (v/v) Glycerol, 0.1% (v/v) Triton X-100, and sonicated on ice with Branson W250 sonifier (Brookfield, CT, USA) until the solution was completely cleared. The cell lysate was centrifuged at 4 °C at 35000 rpm for 1 hour by Beckman Optima™ L-80XP ultracentrifuge equipped with a Ti-45 rotor (Beckman, USA). The cell pellet was discarded, and the supernatant containing the soluble protein was filtered through 0.2 μm membrane and loaded onto a Ni-NTA column that was previously equilibrated with 3 column volume of loading buffer containing 200 mM NaCl, 20 mM Tris-HCl pH 7.5, 20 mM Imidazole. Unbound and non-specifically bound proteins were washed by running 5 column volumes of loading buffer through the column. The hexahistidine-tagged *Cb*FDH proteins were eluted with the elution buffer containing 200 mM NaCl, 20 mM Tris-HCl pH 7.5, and 250 mM Imidazole. The eluted proteins were dialyzed in a dialysis membrane (3 kDa MWCO) against a buffer containing 200 mM NaCl and 20 mM Tris-HCl pH 7.5 overnight at 4 °C to remove excess imidazole. The concentrated pure proteins were run on an 15% SDS-PAGE for verification.

The protein crystallization screening experiments were performed using the sitting-drop microbatch under oil method. The purified wild-type and Val120Thr mutant CbFDH protein were mixed with ∼3500 commercially available sparse matrix crystallization screening conditions in a 1:1 volumetric ratio in 72-Terasaki plates (Greiner Bio-One, Kremsmünster, Austria). The mixtures were covered with 16.6 Μl 100% paraffin oil (Tekkim Kimya, Istanbul, Türkiye). The crystallization plates were incubated at 4 °C and frequently checked under a light microscope. The best CbFDH crystals were grown within six weeks in Wizard III condition #47 (Hampton Research, USA) for both wild-type and mutant protein. This condition consists of 30% (w/v) PEG 5000 MME, 100 mM MES/Sodium hydroxide pH 6.5, and 200 mM Ammonium sulfate.

### 4.2 Activity assays for enzyme kinetics

Activity assays were performed for the wild-type and mutant CbFDH at different substrate, coenzyme, and substrate concentrations in order to calculate the K_m_, V_max_, and k_cat_ values. The values were used to plot the Michael-Menten and Lineweaver-Burk graphs, which yielded an equation for calculations. The reaction conditions used for measurements are shown in Table 3.

**Table 3.** Reaction conditions for activity assays.

| Condition | Concentrations |
| --- | --- |
| Formate concentration (mM) | 0.612, 1.25, 2.50, 5.00, 10.00, 20.00, 40.00 |
| NAD<sup>+</sup> concentration (mM) | 1.00, 2.00, 3.00, 4.00, 5.00, 6.00 |
| *CbFDH* concentration (mg/ml) | 0.0306, 0.0612, 0.125, 0.25, 0.50, 0.75 |

The measurements were performed at 25°C and 340 nm in a microplate reader (Varioscan, ThermoFisher Scientific, USA).

### 4.3 Crystal harvesting and delivery

The *Cb*FDH crystals were harvested by using MiTeGen MicroLoops mounted to a magnetic wand (Garman & Owen, 2006) while simultaneously monitoring under a light microscope, as described by Atalay et al. (2023). The harvested crystals were flash-frozen by plunging them in liquid nitrogen and placed in a cryo-cooled robotic sample puck (Cat#M-CP-111-021, MiTeGen, USA). The puck was mounted to a puck transfer and mounting tool and placed into the sample dewar which is auto refilled with liquid nitrogen at 100 °K, of the *Turkish DeLight* XRD device (Gul et al., 2023).

### 4.4 Data collection and processing

Diffraction data from the *Cb*FDH crystals were collected using Rigaku’s XtaLAB Synergy Flow XRD at University of Health Sciences (Istanbul, Türkiye) controlled with the CrysAlis Pro software version 1.171.42.35a (Rigaku Oxford Diffraction 2021). The crystals were constantly cooled by the Oxford Cryosystem’s Cryostream 800 Plus set to 100 °K. The PhotonJet-R X-ray generator was operated at 40 kV, 30 mA, 1200.0 W with a beam intensity of 10%. The data was collected with 1.54 Å wavelength and the detector distance was 47.00 mm. The scan width was set to 0.25 degrees while exposure time was 10.0 minutes. Data collection was performed for 15 hours 43 minutes 19 seconds.

Data collection at ambient temperature was performed as described by Gul et al. (2023). Data reduction was performed using the CrysAlis Pro version 1.171.42.35a. The data reduction result was obtained as an *.mtz file.

### 4.5 Structure determination and refinement

The cryogenic *Cb*FDH structure was determined at 1.4 Å with the space group P1 using the automated molecular replacement program *PHASER* version 2.8.3 (McCoy et al. 2007) within the *PHENIX* suite version 1.20.1 (Adams et al. 2010). The previously released *Cb*FDH structure with PDB ID: 5DNA was used as an initial search model for molecular replacement of wild-type *CbFDH* structures (Guo et al. 2016). For Val120Thr-*CbFDH*, Val120 of the wild-type *CbFDH* structure was mutated to Thr on PyMoL to be used as the search model in molecular replacement. Initially rigid-body and simulated-annealing refinements were performed by *PHENIX*. Then, individual coordinates and translation/libration/screw (TLS) parameters were refined along with simulated annealing map refinement. The final model of the CbFDH structure was checked by using *COOT* version 0.9.8.1 (Emsley & Cowtan 2004) after each refinement. Water molecules were added into appropriate electron density clouds while those located outside the density were manually removed. The structures were refined until R_work_ and R_free_ values were sufficiently low (Brünger, 1992). The structure figures were generated by using *PyMOL* (Schrödinger, LLC) version 2.5.2 and COOT version 0.9.8.1.

## Supplementary Material

Supplementary Figures.

## Author Contributions

M.G., B.Y., H.B., and H.D. designed the experiment. M.G. and B.Y. performed the protein expression, purification, and crystallization. M.G. and B.Y. performed the data collection, processing, and structure refinement. M.G., B.Y., H.B., and H.D. prepared the manuscript. All authors have read and agreed to the published version of the manuscript.

## Data Availability Statement

The cryogenic wild-type, the ambient temperature wild-type, and the cryogenic Val120Thr mutant CbFDH structures presented in this manuscript have been deposited to the Protein Data Bank under the accession number 8HTY, 8IQ7, and 8IVJ, respectively. Any remaining information can be obtained from the corresponding author upon request.

## Acknowledgments

Authors would like to dedicate this manuscript to the memory of Dr. Albert E. Dahlberg and Dr. Nizar Turker. The authors gratefully acknowledge use of the services and Turkish Light Source (Turkish DeLight) X-ray facility at Sağlik Bilimleri University Deneysel Tip Araştirma ve Uygulama Merkezi (SBU-DETUAM). The authors also gratefully acknowledge use of the services and facilities at Koç University Isbank Research Centre for Infectious Diseases (KUIS-CID).

