## Supplementary Figures for "Structural Analysis of Wild-type and Val120Thr Mutant *Candida boidinii* Formate Dehydrogenase by X-ray Crystallography"

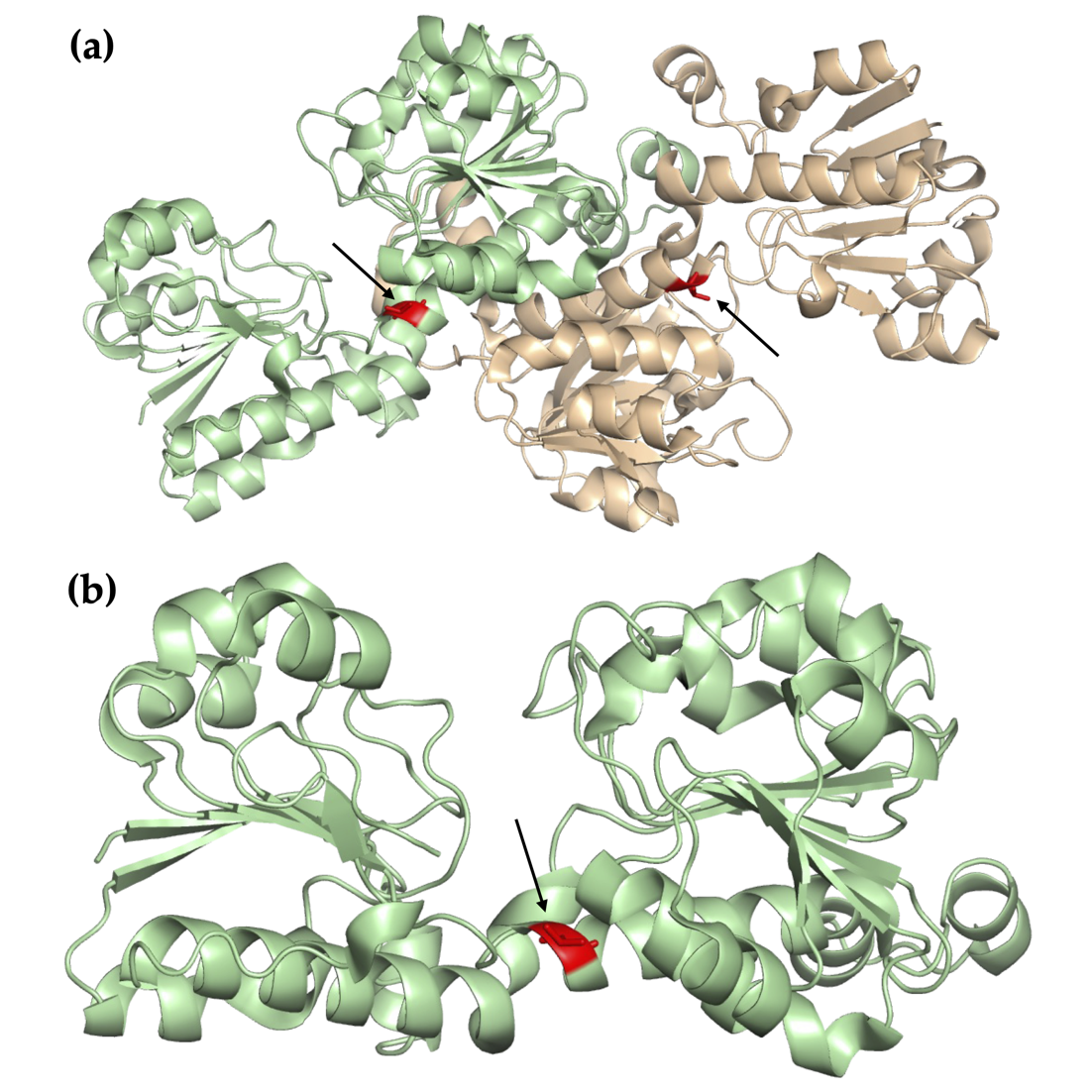

**Supplementary Figure 1.** Position of the 120^th^ residue within *Cb*FDH. **(a)** 120^th^ residues in *Cb*FDH homodimer structure are indicated with red color and arrows. **(b)** 120^th^ residue in *Cb*FDH monomer is indicated with red color and arrow.

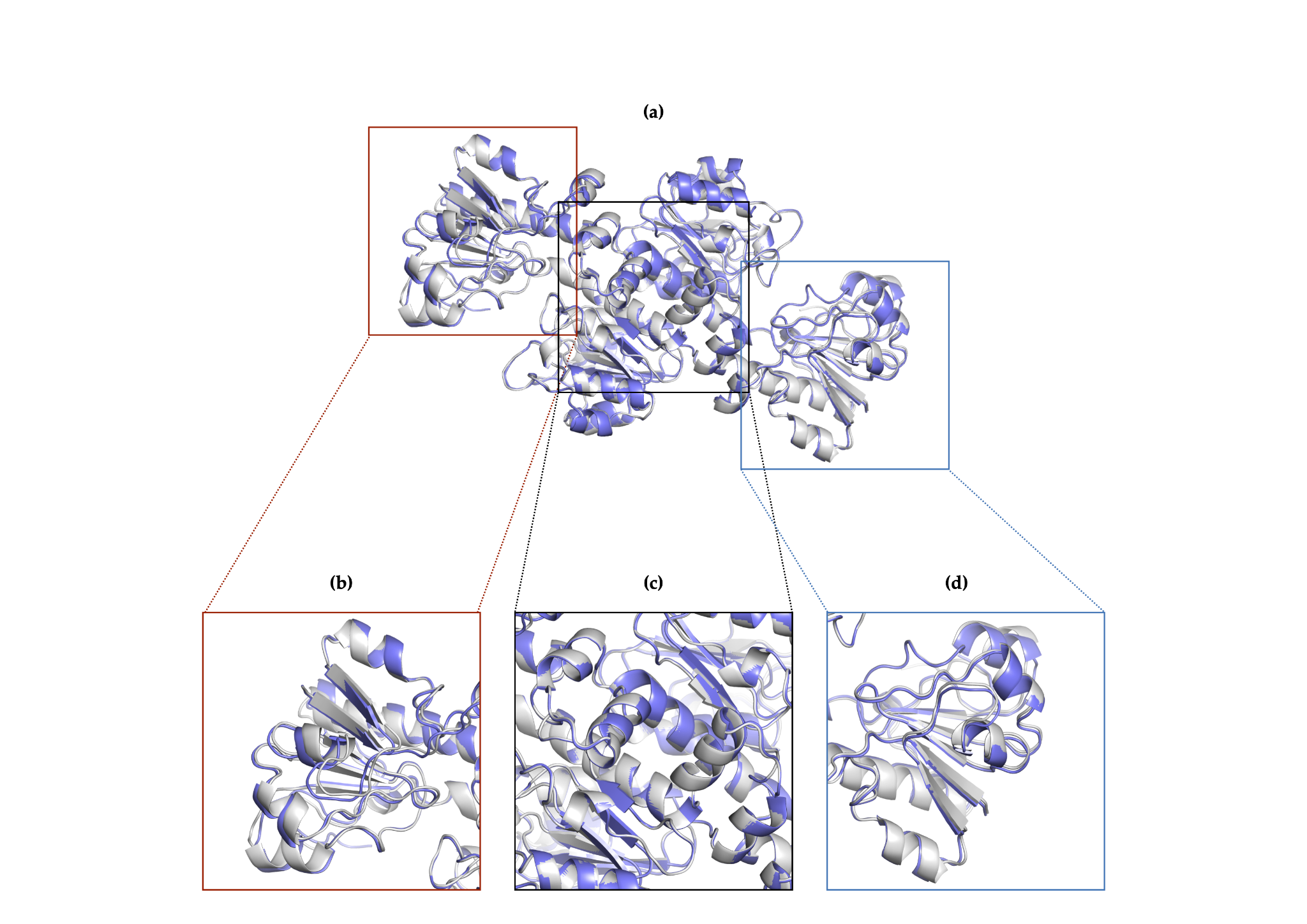

**Supplementary Figure 2.** Superposition of two homodimers of *Cb*FDH. **(a)** Two homodimers superposed. The first homodimer is colored in slate, and the second one is colored in gray. **(b),(d)** Regions that do not participate in dimerization. **(c)** Dimerization region.

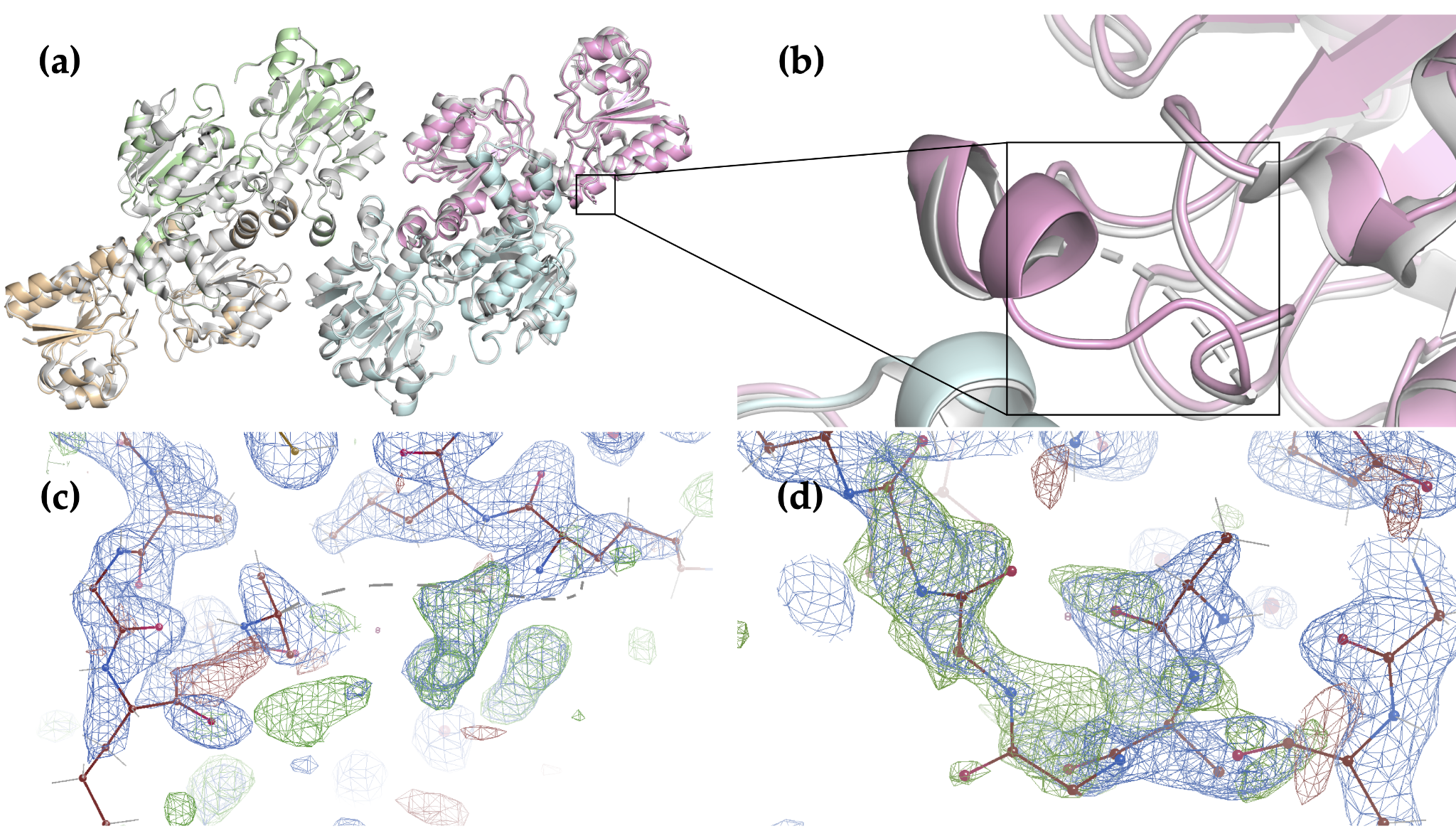

**Supplementary Figure 3.** Chain C residues 15-18 comparison between 8HTY and 5DNA. (**a**) Superposition of 8HTY with 5DNA. The black square indicates the residues 15-18. 8HTY is colored in four colors: pale green, wheat, pink, pale cyan. 5DNA is colored in gray. (**b**) A closer look at the residues 15-18 in the black square. The dashed lines represent the missing residues of 5DNA. (**c**) 2Fo-Fc (colored in slate) and Fo-Fc (colored in green) maps showing the 5DNA chain C residues 14 and 19 and dashed lines for the missing residues. (**d**) 2Fo-Fc and Fo-Fc maps colored same is panel C showing the chain C residues 14-19 belonging to 8HTY.

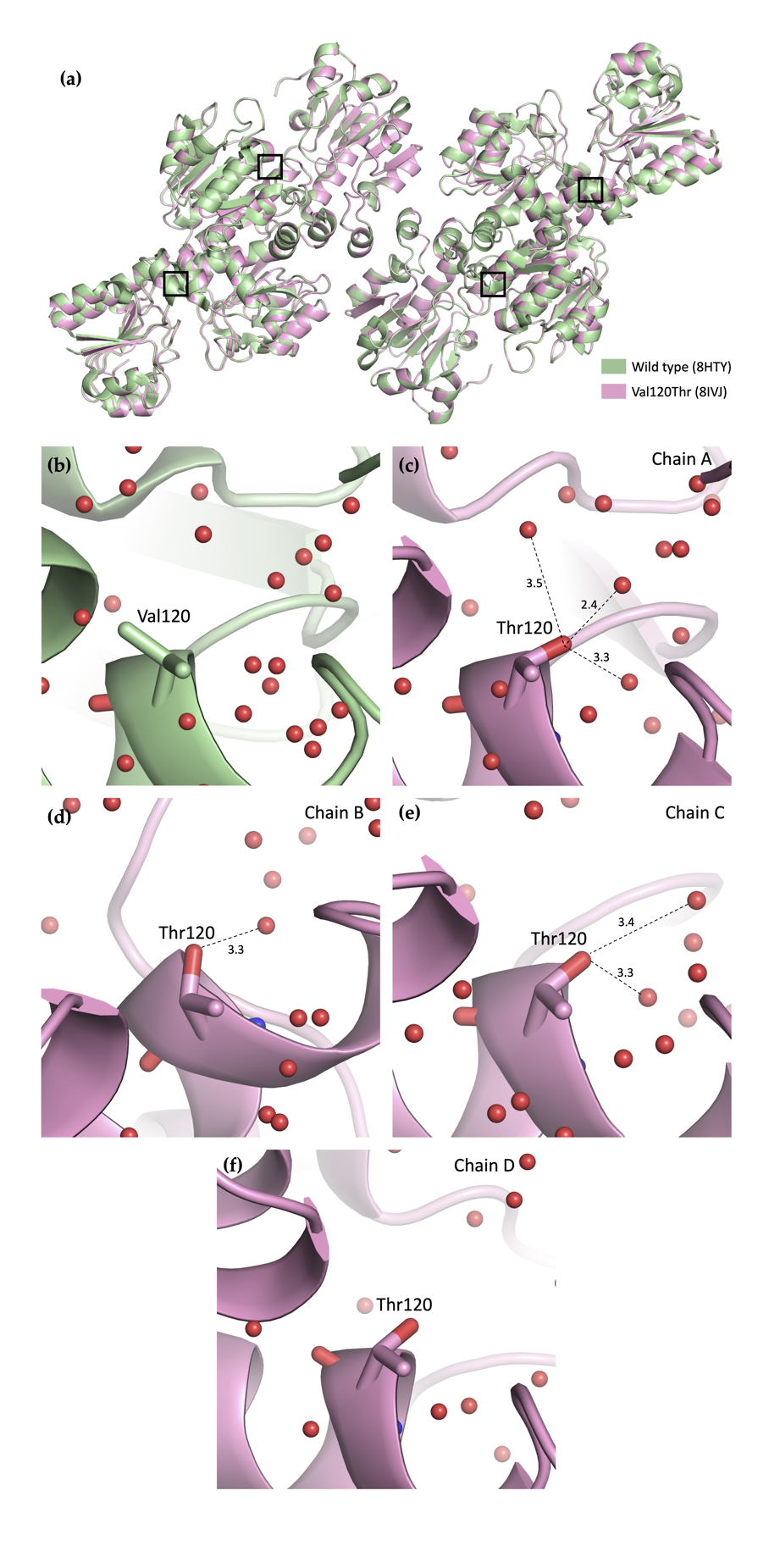

**Supplementary Figure 4.** Comparison of wild-type (PDB ID: 8HTY) and Val120Thr mutant (PDB ID: 8IVJ) *Cb*FDH. **(a)** Superposition of wild-type and Val120Thr mutant *Cb*FDH. Wild-type is shown in pale green, mutant is shown in pink. **(b)** Closer look at the wild-type FDH Val120. **(c, d, e, f)** Closer look at the Val120Thr mutant, showing new hydrogen bonds between Thr120 and water molecules. Hydrogen bonding distances in Angstroms are indicated with dashed lines. Chain ID is indicated in the upper right corner of each panel.
